# A new fitness indicator in birdsong: a trade-off between song length and note length

**DOI:** 10.64898/2025.12.05.692489

**Authors:** Javier Sierro, Selvino R. de Kort, Sue Anne Zollinger, Ian R. Hartley

## Abstract

In acoustic, visual or vibrational displays, motor performance is likely to convey key information on whole-organism condition. To understand the communicative value of such behavioural displays, we must consider the biomechanical trade-offs constraining motor performance. Using song recordings of 140 colour-ringed male and female blue tits (*Cyanistes caeruleus*), we describe a trade-off between song length and note length that defines an upper boundary, possibly arising from respiratory constraints. We measured how close each song lies to this boundary and named this metric song effort score. In males, song effort score peaked during the fertile period and was positively associated with clutch size, describing individual differences in vocal performance relevant to fitness. Recovery score, the analogous metric for the trade-off between note length and gap length, also peaked during the fertile period but was not associated with clutch size. Voiced and unvoiced modulation scores, measures of frequency modulation performance in relation to boundary trade-offs (i.e., vocal deviation), declined or did not vary over the season and were unrelated to clutch size. Across behavioural contexts, song effort score was lower during singing duels and higher during alarm, whereas trills were faster in both contexts, supporting trill rate as an indicator of motivation or arousal. Describing how display components constrain each other can improve the quantification of the individual variation on which selection acts. Ultimately, this approach can help unify our understanding of the role of motor performance in animal communication across display modalities and taxa.

## INTRODUCTION

From drumming woodpeckers (Miles et al. 2018b) to dancing flies (Butterworth et al. 2019; Ishihara and Miyatake 2021), many animals perform motor displays to interact and communicate with other individuals (Anderson et al. 2021; Barske et al. 2011; Clark et al. 2015; Lane and Briffa 2020; Mappes et al. 1996; Nakajima and Onikura 2016). In recent years, behavioural research has converged on the idea that motor performance during animal displays reflects whole-organism condition (Fusani et al. 2014; Lane and Briffa 2021; Schwark et al. 2022) and variation in motor performance can be used to assess individual quality or motivation during both intra- and inter-sexual interactions (Barske et al. 2011; Gilbert and Uetz 2016; Preininger et al. 2013). Motor displays can usually be quantified in several dimensions, and often involve a series of coordinated moves. Almost inevitably, increasing performance in one dimension, such as running speed or calling rate, comes at the cost of decreasing another such as running time or call duration (e.g., Reichert and Gerhardt, 2012). Thus, quantifying only parameters like speed or rates is not adequate to fully assess the performance level of the motor display (Hogan and Stoddard 2018; Reichert and Gerhardt 2012). Understanding such trade-offs is key to establish how animal displays may reflect honest, individual differences in organism condition or quality (Briffa and Sneddon 2007). Physiological or biomechanical limitations underlying performance constraints have been demonstrated in some cases (Miles et al. 2018b; Mowles et al. 2009; Zollinger and Suthers 2004) and hypothesized in others (Podos 1997; Wagner et al. 2012). Furthermore, recent studies indicate that skewed distributions in the phenotypic space towards the boundary trade-off arise from positive selection pressures pushing the performance towards the limit (Logue and Bonnell 2023).

Birdsong performance involves a skilful and rapid coordination of various muscle groups, including superfast muscles (Elemans et al. 2008; Goller 2021). The elaborate phonation mechanisms, as well as the optimal motor patterns used by songbirds (Adam and Elemans 2019; Zollinger and Suthers 2004), strongly suggest that motor performance is under directional selection (Botero and de Kort 2013; Sakata and Vehrencamp 2012). For instance, vocal consistency, a measure of vocal control skill, is a quality signal in some species, associated with higher reproductive success (Botero et al. 2009; Byers et al. 2015; Ferreira et al. 2016; Sierro et al. 2023b) or age (de Kort et al. 2009). Several boundary trade-offs have been described in birdsong, and theoretically, the songs that come closest to this edge reveal what a singer is capable of. Hence, metrics assessing how close a bird sings relative to the boundary are thought to be indicators of performance level in birdsong (Miles et al. 2018b; Podos 1997). Podos (1997) first described one such trade-off between sound bandwidth and note delivery rate, suggesting it was due to a biomechanical limit on frequency modulation speed. More detailed measures have been proposed such as the frequency excursion index (Podos et al. 2016) or (un)voiced frequency modulation score (Dudouit et al. 2022; Geberzahn and Aubin 2014; Vazquez-Cardona et al. 2023), which may better reflect the motor patterns underlying sound production (Riede et al. 2006). In some species, birds are able to discriminate songs based on this limit (Ballentine et al. 2004; Drăgănoiu et al. 2002; Goodwin and Podos 2014; Illes et al. 2006; Phillips and Derryberry 2017a; Phillips and Derryberry 2017b). Despite the evidence for similar trade-offs in many species (Derryberry et al. 2012; Wilson et al. 2014), the exact physiological mechanism underlying the observed trade-off is still unknown. Adjusting the resonance of the vocal tract could be mechanically constrained (Riede et al. 2006), but a recent study found that frequency modulation rates can exceed previously established limitations (Grudens and Islam 2025), in line with the super-fast contraction properties of syringeal muscles (Elemans et al. 2008; Gladman and Elemans 2024).

Another aspect affecting song structure is the capacity to restore the phonatory system in terms of air pressure between subsequent notes using mini-breaths (Hartley 1990; Zollinger and Suthers 2004). In some species, a trade-off exists between note length and gap length (Cardoso and Mota 2007; Logue et al. 2020), which has been related to the process of breathing cycles. The recovery capacity, between notes of varying length, changes daily throughout the dawn chorus (Vazquez-Cardona et al. 2023) but the biological significance of such variation remains unclear. Analogous constraints affecting recovery have been found in woodpecker drumming, where the speed of drumming is limited by body size across species (Miles et al. 2018b).

Salient features are often taken as a measure of individual performance level, such as the peak power or repetition rates. However, it has been shown that individuals must show their capacity to sustain such level of motor performance over a period of time for a robust evaluation of individual capabilities (Bosca et al. 2025; Brandt 2003; de Kort et al. 2009; Mowles et al. 2010; Sierro et al. 2023a). In woodpeckers, the length of a drumming series is associated with stronger sexual selection (Miles et al. 2018b). In some frog species, longer calls are perceived as more attractive (Welch et al. 1998) and in hermit crabs (*Pagurus bernhardus*), longer bouts of rapping result in higher success during fights (Briffa et al. 2003). In lizards, experimentally impaired endurance results in shorter displays, supporting display duration as an honest indication of body condition (Brandt 2003). The biological importance of song length is well known in birds, as song length varies throughout the season (Keating and Reichard 2021; Leitner et al. 2001), is under sexual selection (Eens et al. 1991; Gentner and Hulse 2000; Kempenaers et al. 1997; Mennill et al. 2006; Nolan and Hill 2004), is associated with a better physical condition (Galeotti et al. 1997; Lambrechts and Dhondt 1986) and with higher survival rates (Bijnens 1988). During simulated territorial intrusions, longer songs elicit a stronger response (Lattin and Ritchison 2009; Linhart et al. 2012), although song length is highly variable during agonistic encounters (Sierro et al. 2020). In fact, song overlapping dynamics are key during counter singing (Naguib and Mennill 2010; Poesel and Dabelsteen 2005) and song elongation can result in successfully winning a song duel and thus increased dominance status (Alcami et al. 2025).

Based on previous behavioural (Goto et al. 2025; Podos and Cohn-Haft 2019; Sierro et al. 2023a) and physiological studies (Hartley 1990; Hartley and Suthers 1989), we predicted that a boundary trade-off exists between song length and note length, as we hypothesized that increasing note length will deplete air pressure faster and thus constrain total song length. This is partly the trade-off between call duration and call rate found in frogs (Reichert and Gerhardt 2012), and assessed using a composite metric named *calling effort* (Krueger et al. 2026). Furthermore, as in previous studies, we explored boundary trade-offs between: 1) note length and gap length, 2) note length and frequency travelled and 3) gap length and frequency jump between notes. To understand whether song phenotypic trade-offs provide a reference for birds to signal motivation or quality, we conducted a comprehensive study in male and female blue tit song (*Cyanistes caeruleus*). We recorded 140 colour-ringed males and females repeatedly over three breeding seasons, from three months before laying to incubation, and noted the behavioural context during each recording. In males, we predicted that signals playing a role in gaining mating opportunities should be performed at maximum values during the fertile period (Hau et al. 2017) and be positively associated with reproductive success (i.e., quality signals). Along with quality signals, we predicted that short-term motivational signals would also increase during agonistic singing duels or alarm singing. Repeated observations of the same individuals singing across a range of key moments in their reproductive cycle, including fitness measurements, allow us to draw conclusions on how these sexual signals are modified during inter- and intra-sexual interactions in males. Performance scores are only useful if they add information to the raw traits they combine; hence, we conducted a series of analyses to test how well song metrics explained reproductive success using both the performance scores (i.e., distance to the boundary) as well as the raw song traits constituting the song trade-offs.

## METHODS

### Study population

The study was conducted on a population of free-living blue tits nesting in 110 nest boxes placed in deciduous and mixed woodland on the campus of the University of Lancaster, UK (54.01° N, 2.78° W). This work was part of a long-term study that has been ongoing for over 25 years (Mainwaring and Hartley 2009). Every year, adults were captured at feeding stations using mist nets or trap-door traps in the nest box and colour ringed for identification. Feeding stations were placed for a few days at certain locations and then removed well before the start of the breeding season, to avoid the presence of artificially increased resources (Shutt and Lees 2021). Birds were measured (straightened, flattened, wing length to nearest mm; tarsus length with foot bent down, to nearest 0.1 mm and head-bill length to nearest 0.1 mm), weighed (to nearest 0.1 g) and ringed with a unique combination of three colour rings and one numbered metal ring (Redfern and Clark 2001). During the breeding season, individuals were sexed in the hand based on the presence of a brood patch (females) or cloacal protuberance (males) (Svensson 1992). Birds were also aged as first-year or older, based on plumage characteristics (Svensson 1992). All individuals included in this study were caught at least once during the breeding period. Because otherwise the sex would be uncertain, birds never caught were excluded from the sample. We monitored the nest boxes once every three days during the breeding period to determine occupancy, state of nest building, first egg date and clutch size. The female receptive period, when copulations take place, extends from five days before egg laying through to the completion of the clutch (Kempenaers et al. 1992). In this study, we used clutch size as a proxy for reproductive success (for a discussion on this choice, see (Sierro et al. 2023b)).

Based on structural criteria and on spectrograms reported in previous studies (Bijnens and Dhondt 1984; Mahr et al. 2016), we defined song as a vocalisation composed of a few introductory, higher-frequency notes followed by a trill, which is the last part of the song where a note is repeated in succession (Figure 1) (Bijnens and Dhondt 1984; Cramp and Brooks 1993). A note was defined as a continuous trace on the spectrogram (Knudsen and Gentner 2010). Individual blue tits present song-type repertoires that range from 3 to 8 different types (Bijnens and Dhondt 1984; Sierro et al. 2023b). Song type categorization was made by visual inspection of the spectrograms (Sierro et al. 2023b).

**Figure 1.**
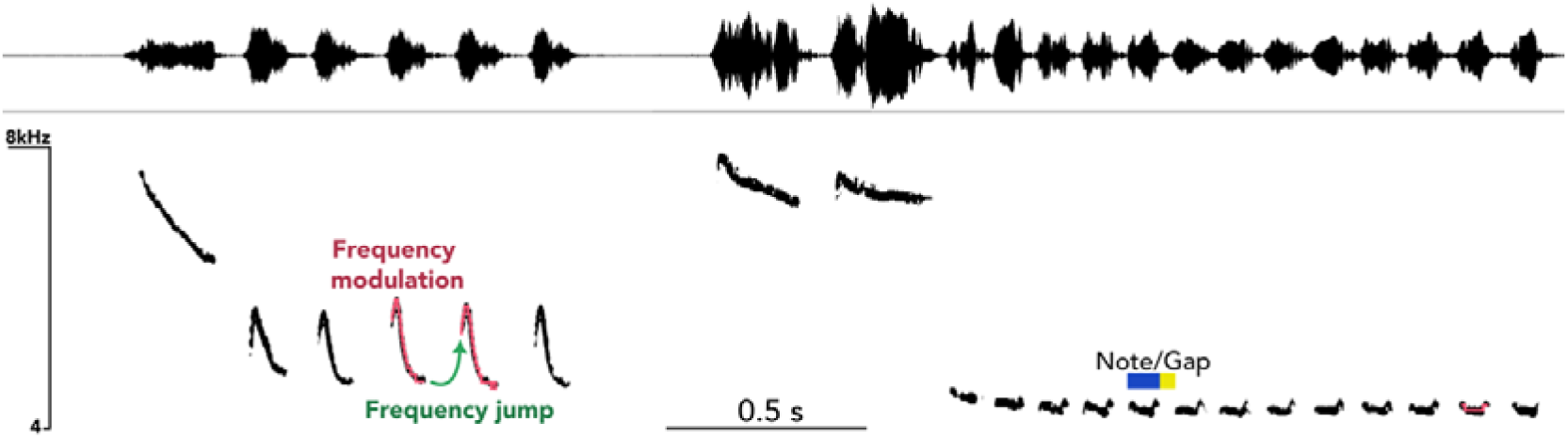
Spectrogram and waveform of two typical blue tit song types from the same individual. The structure includes one or more high-frequency, introductory notes followed by a trill (i.e., the same note repeated multiple times). Pink lines indicate fundamental frequency tracking used to measure frequency modulation travelled. The first and last two points detected in the frequency tracking were used to measure the frequency jump from one note to the next in the same trill (green arrow). Note length (blue box) and gap length (yellow box) are also indicated.

### Ethical statement

All fieldwork involving blue tits was approved by the Lancaster University animal welfare and ethical review board and licensed (BTO ringing license A3673), where appropriate, by Natural England, and the British Trust for Ornithology.

### Audio recordings

In three consecutive years from 2018 to 2020, we recorded songs from adult males and females (Sierro 2022) from January to May. We sampled both sexes to test whether boundary trade-offs constrain birdsong. We recorded songs during walking transects using a Marantz PMD661 recorder (48kHz, 24-bit) and a Sennheiser ME67 microphone. Singing individuals were identified by colour ring combinations. During the winter months of the sampling period (Jan-Mar), fieldwork was conducted during daylight hours, after sunrise until midday. Blue tits increase their singing activity as the season progresses towards the breeding period (April-May), with earlier singing times and longer singing periods (Hinde 1952). We thus gradually advanced our sampling time as the season progressed. Observations were categorised into three distinct behavioural contexts: agonistic interactions, solo songs and alarm. Solo song was characterised by an individual bird singing without any active social interactions with conspecifics or alarm responses. Agonistic interactions were defined by the occurrence of two individuals actively chasing, displacing or attacking each other (Irschick et al. 2007). Finally, alarm context was defined by the presence of a predator and/or observed mobbing behaviour.

### Acoustic analysis

For each song type per recording day and per individual, we selected up to ten songs for acoustic analysis. We analysed each song by manually selecting individual notes on the spectrogram and waveform, using the labelling tool in Audacity to mark the start and end of each note (Mazzoni and Dannenberg 2014). These selections were then fed to a custom-made R script (R Core Team 2022) to extract all relevant song metrics (Ligges 2013; Sueur et al. 2006). We tracked the fundamental frequency of each note based on the *dfreq* function in the ‘seewave’ package (Sueur et al. 2006) (window length = 512, window type = Hamming, 95% overlap). We applied a log10 transformation to the frequency axis (Cardoso 2013) because this resembles how birds perceive sound (Hoeschele et al. 2013; Saunders et al. 1979) and the physical process required to modulate frequency (Fletcher and Rossing 2012). To correct misdetections, we applied an amplitude filter removing frequency detections below 15% of the peak amplitude, normalized within each note, and restricted the frequency modulations to a maximum of 5% of the absolute frequency between consecutive points to avoid tracking of upper harmonics. Finally, we calculated the frequency modulation travelled as the cumulative frequency change along each note. Because the fundamental frequency track is quantized in steps of one FFT bin (93.75 Hz) and contains one point every ∼0.5 ms, summing all point-to-point differences accumulates tracking noise in proportion to note duration. We therefore smoothed each track with a 3-ms running median and summed only the rises and falls between successive turning points that exceeded 1.5 bins (cf. frequency excursion; Podos et al. 2016).

### Statistical analysis

#### Boundary trade-off metrics

We assessed four boundary trade-offs: 1) song length vs. note length, 2) note length vs. gap length, 3) frequency travelled vs. note length and 4) frequency jump between notes vs. gap length. Because neither trait in a pair can be considered the cause of the trade-off, each boundary was estimated with the reciprocal quantile regression method (Cardoso 2019): two mixed-effects quantile regressions (‘lqmm‘ package (Geraci 2014)), each with one trait of the pair as the response, individual identity as a random intercept, and sex and its interaction with the predictor to obtain sex-specific boundaries. The boundary trade-off was the bisector of the two reciprocal lines, estimated on z-standardized traits so that both traits weighed equally. A trade-off was considered present when the 95% CI of the boundary slope did not include zero (Gelman and Hill 2007). For each observation, we then measured its signed orthogonal distance to the boundary: a value of zero means that a song (or note) lies on the boundary, and increasingly negative values mean that it lies further inside the range of songs that all birds produce. Note that the boundary is not a hard limit as we do not expect individuals to be singing at maximal intensity. Thus, values can lie beyond the boundary as it is only a reference to assess the position in the boundary trade-off, not the absolute limit in performance. We also compared this metric to the consensus measure proposed by Cardoso (Cardoso 2019) and found them to be nearly identical, as predicted by theory. See Supplementary Methods for more details.

For song length-note length boundary analyses, we averaged note length per song and then calculated the proximity of each song to the boundary. This variable was named *song effort score,* following the terminology proposed for comparable trade-offs in frogs (Krueger et al. 2026; Reichert and Gerhardt 2012). Recovery score was the distance to the note length-gap length boundary, voiced modulation score was the distance to the frequency travelled-note length boundary and unvoiced modulation score was the distance to the frequency jump-gap length boundary trade-off.

#### Seasonal and contextual variation of song

To determine whether individuals modified their songs throughout the season and with context, we restricted our sample to individuals that had been recorded singing the same song type on at least three different dates along the season. We pooled together three years of recordings and for each individual and song type, all metrics were mean centred and scaled by dividing by one standard deviation, again within individual and song type. This removes all differences between individuals and between song types, so that the models describe only how each individual changed his own songs over the season and between contexts. This method results in normalized song variables that can be compared, where positive values indicate individual values that are higher than the mean. Furthermore, this approach allowed us to understand whether song structure varied in relation to behavioural context, comparing song variation within individual and song type while controlling for seasonal changes. Using normalized variables of song effort score, recovery score, voiced modulation score and unvoiced modulation score, we fitted four separate generalized additive mixed-effects models (GAMM). For females, our sample of individuals recorded singing the same song type at three different dates was relatively small (N = 11, 22.0 ± 13.5 songs per individual) and often during a single context category, compared with males (N = 68, 76.4 ± 57.8 songs per individual). Thus, we restricted the analysis of seasonal and contextual variation to males only. We restricted the period of analysis to those weeks where more than three individuals were recorded. For each model, one song metric was the response variable, as a function of time, measured as weeks from the first egg for each male (Schlicht and Kempenaers 2020). We used GAMs because they are suitable to analyse non-linear variation over time series (smooth term) while estimating the linear parameters of other variables (parametric term) (Zuur et al. 2009). The seasonal variable was weeks to first egg, defined as the number of weeks before (negative) or after (positive) the date of first egg in the nest of each individual (date of first egg = 0). Hence, this variable was standardized for all three years, regardless of between-year variation in breeding dates. In the models, we included the weeks to first egg in the smooth term. The amount of smoothing was estimated by restricted maximum likelihood (REML), using cubic regression splines with a basis dimension of five. Behavioural context (solo song, agonistic or alarm), was included as a fixed effect in the parametric term. For a robust interpretation of variation in the trade-off metrics, we also present the seasonal variation of the song variables which make up each trade-off, namely, song length, note length, gap length, frequency travelled, frequency jump as well as in trill rate, which is a variable traditionally included in song performance constraints studies and typically found to convey arousal (Linhart et al. 2013). In all cases, variables were normalized within individual and song type as explained before. Test statistics are derived from the frequentist properties of Bayesian confidence intervals for smooth functions (Marra and Wood 2012). The individual’s identity was explicitly included as a random effect smoothing term to control for the repeated measurements design and avoid pseudo-replication.

#### Song and fitness variation

To investigate the relationship between song metrics and fitness, we first calculated a mean value of each song performance parameter per individual and per breeding attempt across the three years of fieldwork. Our sample of female song was relatively small per breeding attempt with only 12 females having more than 10 songs recorded. Hence, we restricted this analysis to males only. To evaluate the effect of song traits on clutch size, our proxy for individual reproductive success (Sierro et al. 2023b), we fitted a generalized Poisson Generalized Linear Mixed Model (GLMM) with a log-link function using the ‘glmmTMB’ package (Magnusson et al. 2017). This model structure was selected to account for the bounded, count nature of the response variable while explicitly controlling for potential underdispersion. Clutch size was modelled as a function of song effort score, recovery score, voiced modulation score, unvoiced modulation score, laying date (days after 1^st^ of April), (Klomp 1970) and the age of the male’s breeding partner, a two-level factor (first-year vs. older; first-year as reference). All continuous predictors were scaled (Gelman 2008). Male identity was included as a random intercept to account for repeated breeding attempts by the same male. We subsequently performed multi-model inference using the *dredge* and *model.avg* functions from the ‘MuMIn’ package (Barton 2011), averaging the top-ranked models within ΔAICc < 2 of the best model. Parameter estimates, standard errors and P values were derived from the full model average, with Wald 95% confidence intervals (CI); an effect was considered significant when its 95% CI did not overlap zero. We assessed collinearity among predictors with the variance inflation factor (VIF) in the ‘performance’ package, using a threshold of 5 (Zuur et al. 2009), and validated the model by inspecting simulated quantile residuals with the ‘DHARMa’ package (Hartig and Hartig 2017).

Because each performance score combines two song traits, we ran four additional analyses to interpret the averaged model and to validate the use of performance scores in contrast with raw song traits that constitute the trade-offs (details in Supplementary Methods). First, we fitted each score as the only song predictor to obtain its effect independently of the other scores. Second, we compared the models built with the performance scores and with the raw traits, to test whether the scores were more or less informative. Third, we tested whether our metrics were robust to repertoire composition, indicating general individual capacity, or differential song type usage. Finally, to interpret the partial effect of recovery score (see Results), we re-expressed note length and gap length as two uncorrelated axes: tempo (notes and gaps both long, i.e., slow trills) and contrast (notes long relative to their gaps).

### Data and code availability statement

The data sets and code used in this study are publicly available at [DOI hidden until publication; Data provided with submission].

## RESULTS

### Description of the sample

All measures are presented as mean ± one standard deviation (SD), unless stated otherwise. We analysed 45,575 single notes (419 ± 334 notes per male and 57 ± 72 notes per female) from 8,176 songs (74 ± 57 songs per male and 13 ± 13 songs per female) sung by 140 individual blue tits (104 males and 36 females). Data sets used to estimate performance trade-offs come from the same 140 individuals, but the exact sample of notes varied because 1) the final notes of each song had, by definition, no associated gap and 2) frequency measurements failed in some notes and were removed. Furthermore, *song effort score* was measured per song, not per note unit. Specific sample sizes for each model are specified in Table S1. From a total of 763 individual singing instances, 689 were recorded during solo context, 30 were recorded during agonistic interactions and 44 instances were recorded during alarm context.

### Boundary trade-offs in song

In males, all four boundary trade-offs were significant, whereas in females the song length–note length and note length–gap length boundaries were not, likely because of the smaller sample (Figures 2, S1, S2; Table S1). Boundary slopes did not differ between sexes, and only the position of the song length–note length boundary was slightly higher in males (Table S1). Distances to the bisector were virtually identical to the consensus metric from Cardoso (2019) (r ≥ 0.998; Figure S1).

**Figure 2.**
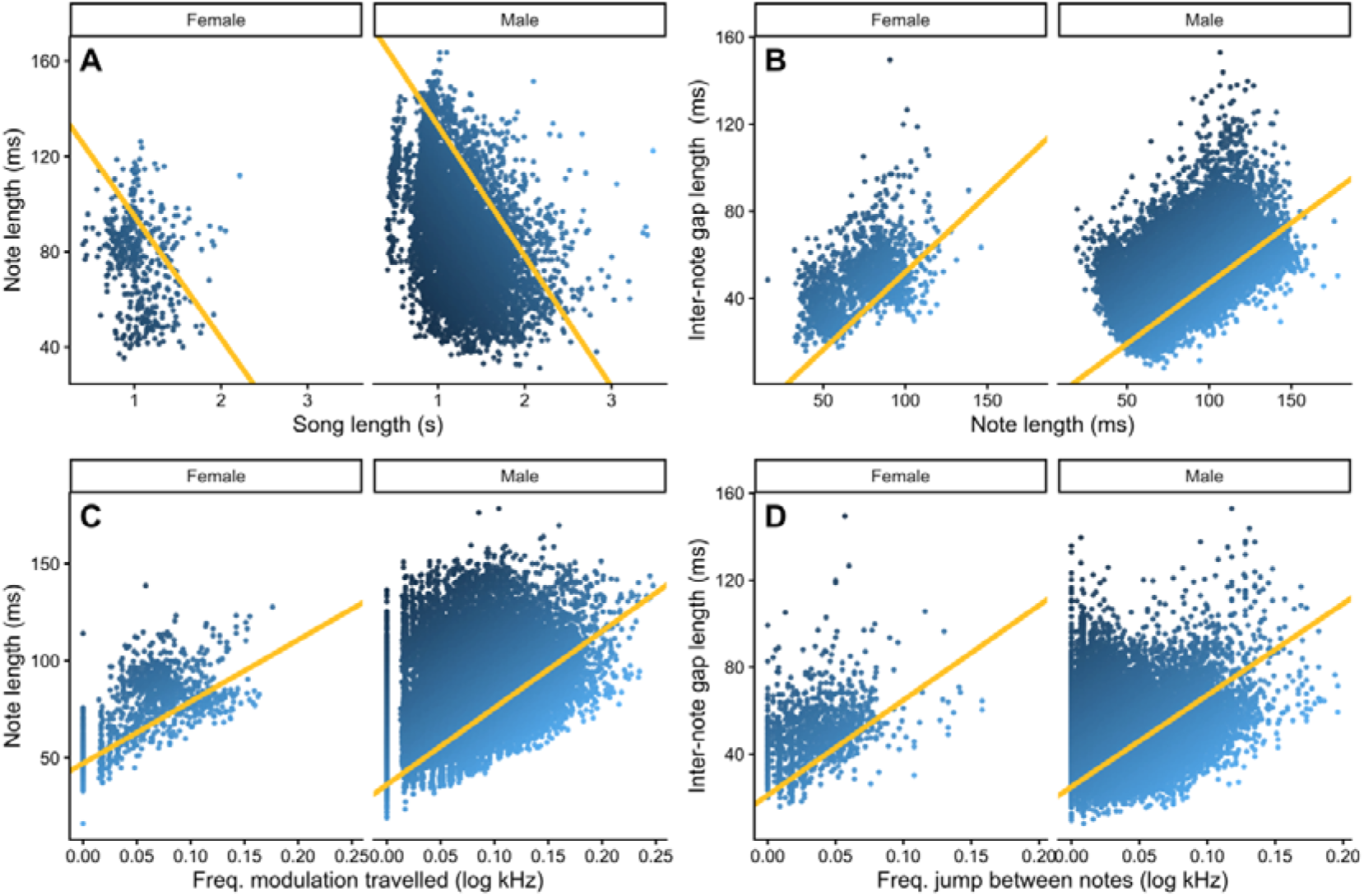
Panel showing the bidimensional phenotypic space described by A) note length and song length, B) gap length and note length, C) note length and frequency travelled and D) gap length and frequency jump between notes. In each case, the estimated boundary trade-off (yellow line) is the bisector of the two reciprocal quantile regression lines (see Figure S1). The blue colour gradient indicates the position of each note (each song in A) as the orthogonal distance to the estimated boundary. Positive values indicate observations beyond the boundary and negative values observations within the phenotypic space.

### Song variation over the season and by context

We found that both the song effort score and the recovery score increased significantly over the breeding season, showing maximum values during the fertile period (Figure 3A-B, Table S2 & S3). This seasonal variation occurred along with changes in the song traits that constitute the trade-offs, namely, song length and note length increased and gap length was lower during the fertile period (Figure 3B, Table S2 & S3). Despite shorter gaps, note length increased so that trill rate was lower during the fertile period (Figure 3B, Table S2 & S3). Voiced modulation score decreased significantly over the season reaching minimum values right after the fertile period (Figure 3C, Table S2 & S3). These changes were likely due to increased note length, since frequency travelled itself did not vary (Figure 3C, Table S3). Unvoiced modulation score showed a nonlinear pattern but the variation associated with season was overall non-significant (Figure 3D, Table S2 & S3). However, frequency jumps declined significantly over the season with low values during the fertile period (Figure 3D, Table S3).

**Figure 3.**
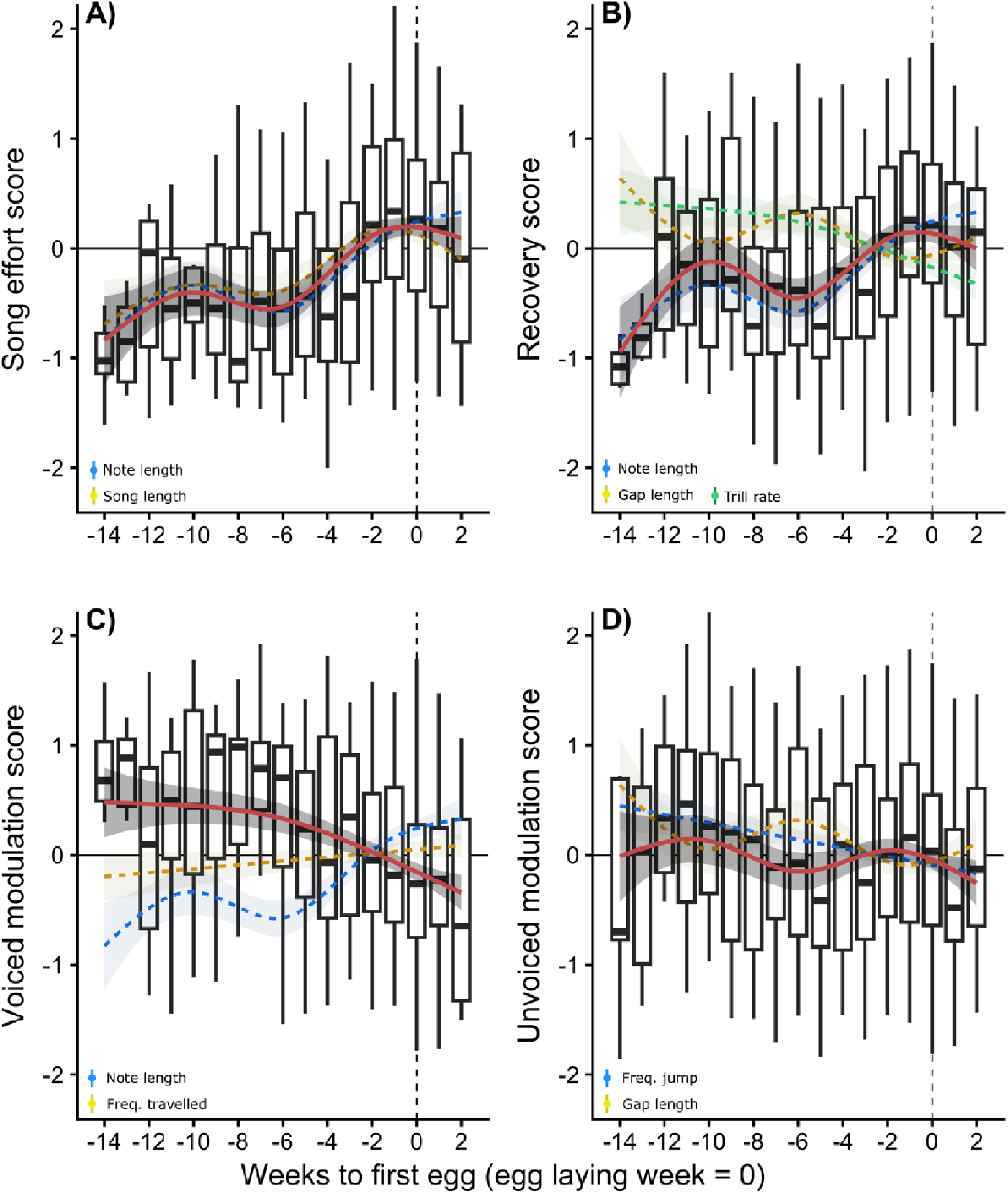
Seasonal variation in male blue tit song. The four trade-off metrics are shown in red in A-D, derived from the boundary trade-offs between two variables, which are shown in the same plots with yellow and blue dashed lines (see legend in each case). Box and whisker plots show median, upper and lower quartiles, and 1.5 interquartile range of all observations within the same week. The shaded area around the lines delimits the estimated 95% confidence interval from the GAM models. Following Kempenaers et al. (1992) the fertile period starts five days before egg laying (first egg week = 0; vertical dashed line). Trill rate, as a signal associated with motivation and performance level, is shown in B (green).

With respect to behavioural context, song effort score was lower during agonistic encounters and higher during alarm than during solo song (Figure 4C; Table 1), mirroring song length, which was lower during agonistic encounters and higher during alarm (Table S4). Recovery score and note length did not change significantly between contexts (Figure 4D; Tables 1 & S4). Gaps were shorter during both agonistic and alarm contexts than during solo song, leading to faster trills (Figure 4D; Table S4). Neither voiced modulation score nor frequency travelled varied with context (Figure 4E; Table 1). Unvoiced modulation score was higher during agonistic encounters (Table 1), reflecting shorter gaps, since frequency jumps did not change with context (Figure 4E–F; Table S4).

**Figure 4.**
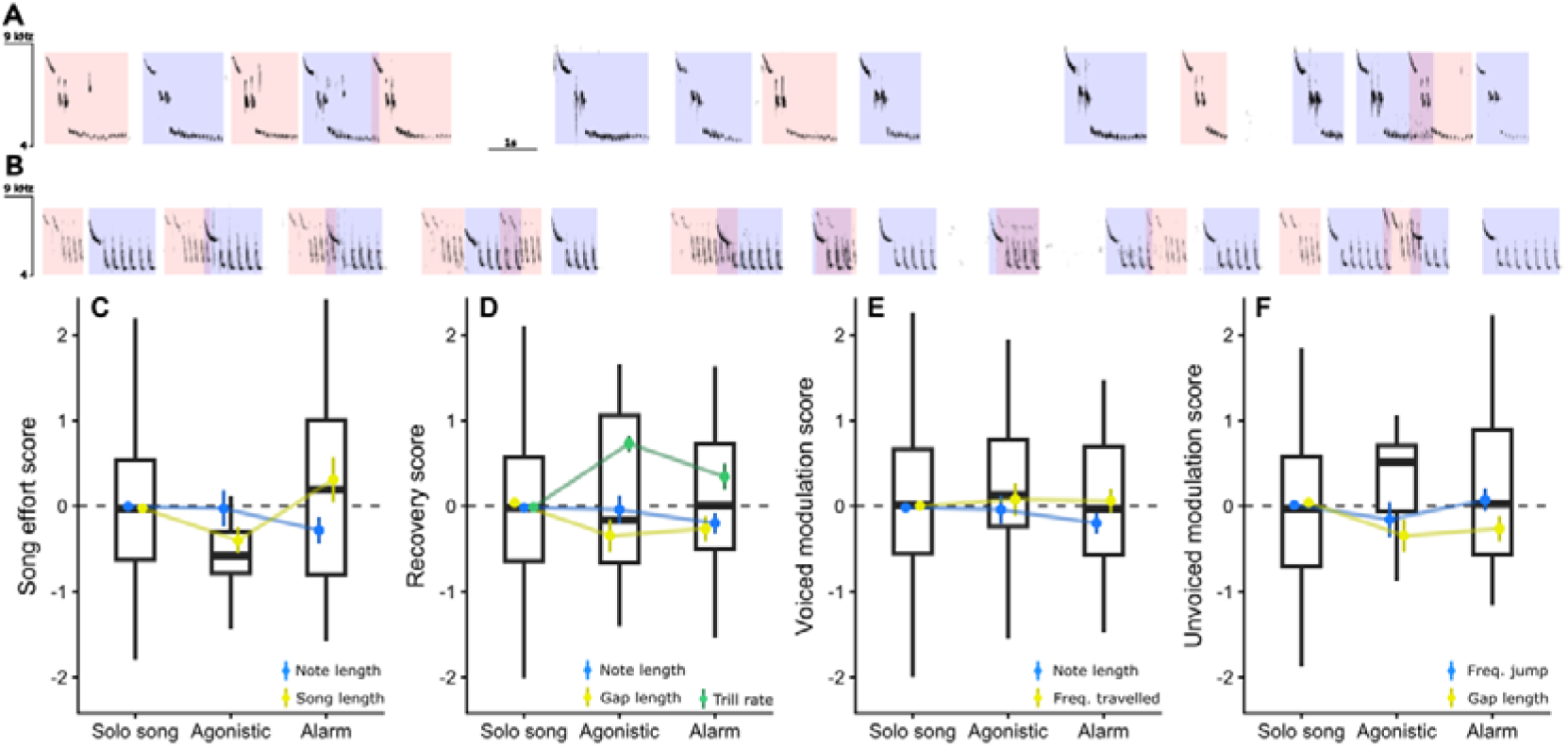
Differences in song structure associated with context. A-B show two examples of blue tit song contests between males. Coloured boxes mark the two individuals to facilitate tracking along the song contest. C-F show the differences in song structure between solo song, agonistic contexts and alarm singing, normalized within individual and song type. Boxplots show the variation in the trade-off derived metrics (i.e., orthogonal distance to the boundary) along with the song trait pairs that constitute the boundary trade-off itself, shown as coloured points and error bars indicating mean ± the SE of the mean. All variables were standardized within individual and song type and adjusted for season (GAMM seasonal smooth removed).

**Table 1.**
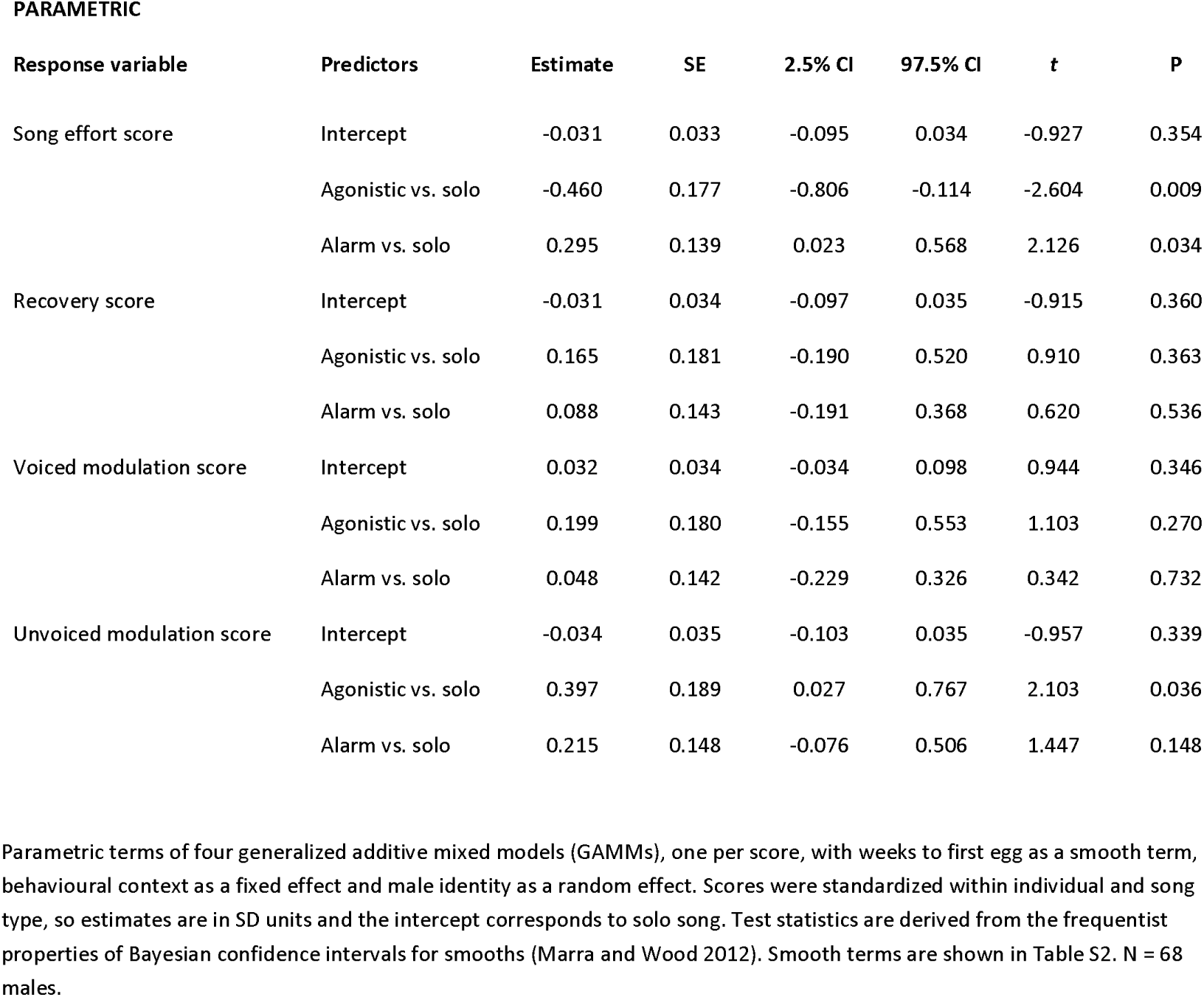
Effect of behavioural context on song performance scores in male blue tits.

| PARAMETRIC |  |  |  |  |  |  |  |
| --- | --- | --- | --- | --- | --- | --- | --- |
| Response variable | Predictors | Estimate | SE | 2.5% CI | 97.5% CI | t | P |
| Song effort score | Intercept | -0.031 | 0.033 | -0.095 | 0.034 | -0.927 | 0.354 |
|  | Agonistic vs. solo | -0.460 | 0.177 | -0.806 | -0.114 | -2.604 | 0.009 |
|  | Alarm vs. solo | 0.295 | 0.139 | 0.023 | 0.568 | 2.126 | 0.034 |
| Recovery score | Intercept | -0.031 | 0.034 | -0.097 | 0.035 | -0.915 | 0.360 |
|  | Agonistic vs. solo | 0.165 | 0.181 | -0.190 | 0.520 | 0.910 | 0.363 |
|  | Alarm vs. solo | 0.088 | 0.143 | -0.191 | 0.368 | 0.620 | 0.536 |
| Voiced modulation score | Intercept | 0.032 | 0.034 | -0.034 | 0.098 | 0.944 | 0.346 |
|  | Agonistic vs. solo | 0.199 | 0.180 | -0.155 | 0.553 | 1.103 | 0.270 |
|  | Alarm vs. solo | 0.048 | 0.142 | -0.229 | 0.326 | 0.342 | 0.732 |
| Unvoiced modulation score | Intercept | -0.034 | 0.035 | -0.103 | 0.035 | -0.957 | 0.339 |
|  | Agonistic vs. solo | 0.397 | 0.189 | 0.027 | 0.767 | 2.103 | 0.036 |
|  | Alarm vs. solo | 0.215 | 0.148 | -0.076 | 0.506 | 1.447 | 0.148 |

### Song and fitness variation

Song effort score was positively associated with clutch size in the averaged model and was present in all selected models (Figure 5, Table 2 & S5). With a 1 SD increase in song effort score, there is a 7.4% increase in clutch size, similar in magnitude and importance to the effect of laying date (−7.9% per SD, Table 2). Song effort score was also the only score associated with clutch size when fitted as the only song predictor (Table S6). Recovery score showed a negative partial effect in the averaged model (Table 2) but was unrelated to clutch size on its own (Table S6). Neither voiced nor unvoiced modulation score was associated with clutch size (Tables 2, S5 & S6). All VIFs were ≤ 3 and residual diagnostics showed no deviations (Figure S3).

**Figure 5.**
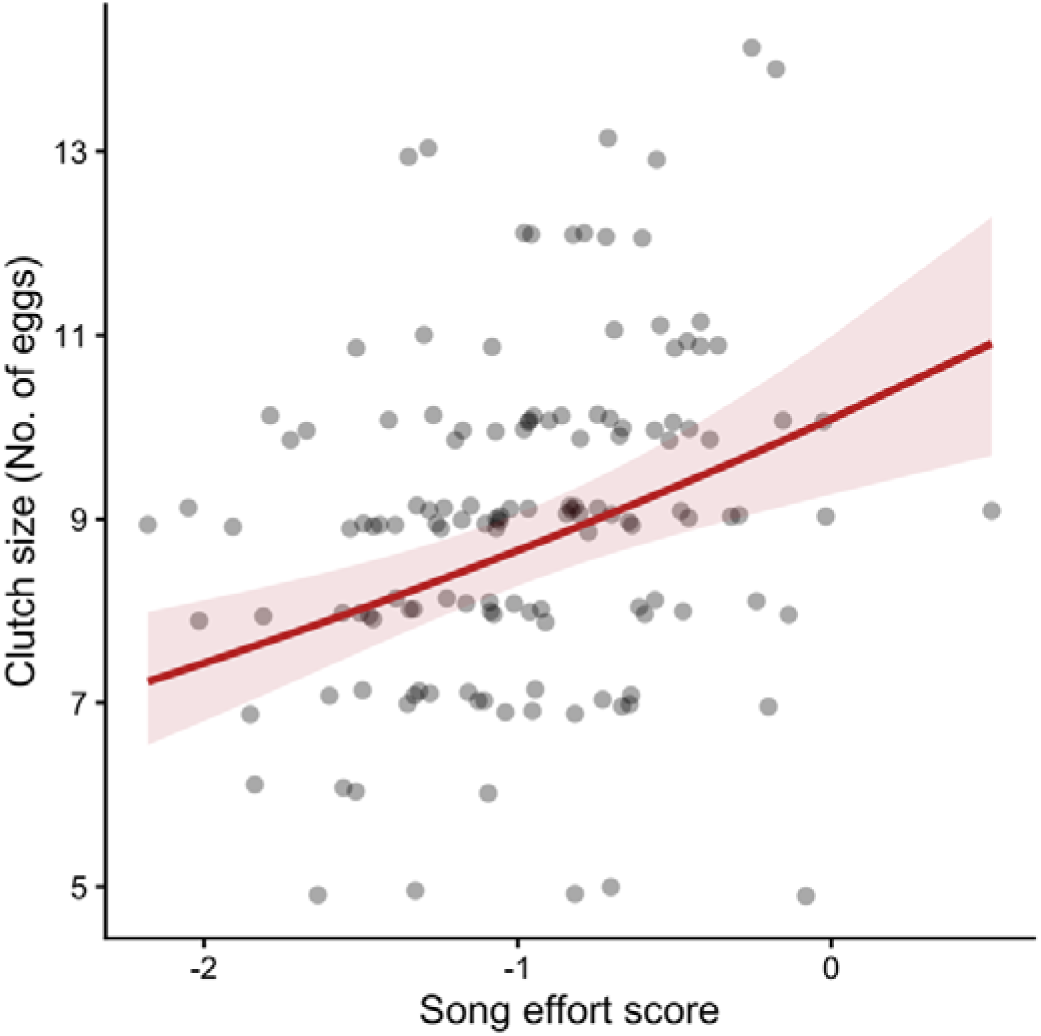
Song effort score and its association with clutch size. The line and shaded band show the predicted clutch size (± 95% CI) from the full model, with the remaining continuous predictors held at their mean, partner age at its reference level (first-year) and the random effect of individual set to zero. Points show the raw data, with transparency and a small artificial displacement of 0.15 in height to allow visualization of overlapping data points.

**Table 2.** Clutch size as a function of song performance scores in male blue tits.

| Variable | Estimate | SE | 2.5% CI | 97.5% CI | Z | P | Relative importance |
| --- | --- | --- | --- | --- | --- | --- | --- |
| Intercept | 2.177 | 0.023 | 2.132 | 2.222 | 95.324 | < 0.001 | - |
| Laying date | -0.082 | 0.013 | -0.108 | -0.057 | -6.24 | < 0.001 | 1 |
| Partner age<br>(Older vs. First year) | 0.017 | 0.028 | -0.038 | 0.072 | 0.619 | 0.536 | 0.416 |
| Song effort score | 0.071 | 0.017 | 0.038 | 0.105 | 4.146 | < 0.001 | 1 |
| Recovery score | -0.052 | 0.018 | -0.089 | -0.016 | -2.849 | 0.004 | 1 |
| Voiced modulation score | 0.001 | 0.007 | -0.012 | 0.014 | 0.177 | 0.860 | 0.116 |
| Unvoiced modulation score | 0.006 | 0.013 | -0.02 | 0.032 | 0.461 | 0.645 | 0.328 |

Full average of the generalized Poisson GLMMs within $\Delta AIC_c < 2$ of the best model, with laying date and partner age (older vs. first-year) as covariates and male identity as a random intercept. All continuous predictors were scaled (SD units); estimates are on the log scale. Relative importance is the sum of Akaike weights of the averaged models containing each term. N = 138 breeding attempts from 96 males.

### Performance scores vs. using raw traits

Song effort score predicted clutch size better than its constituent traits (Table S7, Figure S4). On its own, it improved fit by 7.85 AICc units over the covariates-only model, whereas song length alone did not (ΔAICc = −0.37). Song length was associated with clutch size only when note length was added to the model (Table S7), and this two-trait model still fitted 2.3 AICc units worse than song effort score alone, with one more parameter. When all traits were modelled together, the model with the four performance scores (i.e., distances to the boundary) and the model with the five raw traits fitted the data similarly (ΔAICc = 0.25; Table S7).

Song effort score was positively associated with clutch size both within song types and between song types (Table S8A). Its association was therefore robust to repertoire composition, indicating that song effort score can estimate individual performance regardless of song-type categories. In contrast, raw song length had opposite effects within and between song types (Table S8B), consistent with song types of longer songs having shorter notes (Figure S5), which masks the effect of song length when note length is ignored. Finally, when note length and gap length were expressed as two uncorrelated axes, clutch size increased with tempo (longer notes and longer gaps) but not with contrast (notes long relative to their gaps; Table S8C, Figure S6).

## DISCUSSION

We found a trade-off between song length and note length in males, with a similar but non-significant boundary in female blue tits: the longest songs were sung with short notes, and vice versa. We measured the distance of each song to the boundary (i.e., song effort score) as a potential indicator of performance level. In males, song effort score peaked during the female fertile period and was associated with larger clutches. This association went beyond what song length alone could explain and was not affected by song-type categories. By contrast, during agonistic encounters males sang shorter songs and faster trills, with an overall lower song effort score. Recovery score also peaked during the fertile period but was not associated with clutch size or with context. Voiced modulation score declined as the fertile period approached, did not explain variation in clutch size and did not change with context. Unvoiced modulation scores showed no seasonal variation, no association with clutch size and an increase during agonistic interactions. However, this change with context was fully explained by shorter gaps alone (i.e., trill rate). Hence our findings suggest that frequency modulation scores play a minor role in blue tit song communication, with no clear link to fertility or reproductive success. Together, these results are consistent with positive selection on song effort score, taking clutch size as a proxy for reproductive success, and a distinct role of trill rate in aggressive and alarm contexts (i.e., high arousal). This newly described trade-off thus provides a robust metric of individual vocal performance capacity despite the complex interplay of performance constraints and song repertoires.

### *Song effort score*: a boundary trade-off relevant in communication

We found a significant trade-off between song length and note length on the upper boundary of the phenotypic space. Namely, longer songs near this boundary were associated with shorter notes or reversely, longer notes were associated with shorter songs. We hypothesize that longer notes deplete sub-syringeal pressure faster (Goto et al. 2025; Podos and Cohn-Haft 2019) and this may limit song length. Thus, we suggest that song effort score reflects whole-organism condition in birdsong associated with motor control and respiratory capacity (Goto et al. 2025; Hartley and Suthers 1989; Podos and Cohn-Haft 2019). An intriguing finding is that this trade-off explains the adherence to Menzerath’s law observed in many species, e.g., (Gustison et al. 2016; Zhang et al. 2024), the tendency for longer phrases to consist of shorter constituent parts (Menzereth 1954). While the pattern found in blue tits fits this phenomenon, the sequences analysed consist of repeating notes with seemingly equal information content. This supports the biomechanical, perhaps respiratory, constraint as the underlying mechanistic explanation rather than an information-theoretic optimization rule.

Several studies have found longer songs to be sexually attractive (Gentner and Hulse 2000), associated with increased paternity (Eens et al. 1991; Garrido Coria et al. 2025; Kempenaers et al. 1997), and relevant during aggressive encounters (Nelson and Poesel 2012; Sierro et al. 2023a). Our results suggest that the information carried by song length may only be fully revealed relative to the trade-off with note length. Across songbird species (Sierro et al. 2023a) and within individuals (Bosca et al. 2025), vocal performance declines when individuals produce longer songs, supporting song effort score as a relevant parameter beyond blue tits. In fact, studies in frogs (*Hyla versicolor* and *Hyla chrysoscelis*) show a similar trade-off in vocal performance (Reichert and Gerhardt 2012) and females assess vocal display as an integrative performance, not as the value of raw traits in the trade-off (Krueger et al. 2026). From fiddler crabs to frogs or songbirds, multiple components of motor displays are interdependent, constrained by shared physiological or biomechanical limits (Brandt 2003; Miles et al. 2018a; Mowles et al. 2017), and receivers may assess the integrated output rather than its separate components (Krueger et al. 2026; Sierro et al. 2023a; Ward et al. 2013).

### Biological significance of song metrics based on phenotypic trade-offs

The individual decisions and social interactions taking place during the fertile window will have a profound impact on individual fitness (Hau et al. 2017). Specifically, individuals able to gain more within- and extra-pair copulations, reduce cuckoldry and maximize their partner’s investment will have a reproductive advantage. Hence, song signals used to modulate such interactions should present a temporal synchrony with the fertile period of females (Hau et al. 2017), as found for song effort score and recovery score. Physiological changes associated with practice (Adam et al. 2023), neurogenesis and hormonal effects (Caro et al. 2005; Smith et al. 1997) could result in seasonal differences associated with individual attributes. We observed how both song length and note length increased seasonally, so that the changes in song effort score were associated with maximizing both traits in the trade-off. As males singing with higher song effort score also had larger clutches, the evidence supports this trait as a fitness indicator in blue tits. Clutch size also reflects female investment, hence this association could arise from males with higher song effort score being paired with higher-quality females (assortative mating) (Jiang et al. 2013) or from higher female investment (differential allocation) (Sheldon 2000). In either case, song effort score would carry information relevant to reproductive success although demonstrating positive selection on song effort score will require measures of realized reproductive success, such as paternity or recruitment.

Song length by itself was a poor predictor of reproductive success: its effect appeared only once note length was accounted for, and it had opposite effects within and between song types. Namely, within song types, males singing longer songs had larger clutches, whereas males using longer song types had smaller clutches, likely due to a confounding effect of note length and song types. Song effort score, instead, was positively associated with clutch size both within and between song types. That is, males with larger clutches sang song types closer to the boundary and, within each song type, these males sang closer to the boundary than other males using the same types. Song-type categories may therefore partly reflect differences in performance level, depending on how song types are defined in each study system. Ignoring this trade-off could result in a biased assessment of individual performance due to repertoire composition, through differential song-type use or uneven sampling of repertoires. Song effort score, by contrast, captured individual differences relevant to fitness regardless of repertoire composition.

Contrary to the association with female fertility and male fitness, we found that song effort score decreased during agonistic encounters, as males sang shorter songs with high trill rates. This could be explained by the dynamic nature of singing duels where song matching and overlapping are key signals (Akcay et al. 2013; Ballentine et al. 2008), dictating the song rates and song length to modulate conflict (de)escalation (Arnott and Elwood 2009). Nevertheless, males with a higher song effort score may still have an advantage in song duels as eventually elongating some songs could lead to higher dominance (Alcami et al. 2025). During alarm singing, maximizing song effort score could result in higher communication efficiency as longer songs increase signal transmission and detection. The increase in trill rate during alarm further supports this trait as a dynamic parameter related to motivation or arousal (Funghi et al. 2015; Linhart et al. 2013).

We also confirmed a boundary trade-off between note length and gap length (Cardoso et al. 2007; Logue et al. 2020; Mota and Cardoso 2001), which is thought to result from use of mini-breaths to replenish air pressure in the respiratory organs (Hartley 1990). Males modulated their songs and performed with peak recovery score during the female fertile period. Although recovery score was unrelated to clutch size on its own, it showed a negative partial effect once song effort score was accounted, likely due to a confounding effect in the correlation between note length and gap length. Re-expressing note and gap length as two uncorrelated axes showed that clutch size increased with overall longer notes, and gaps, but not with longer gaps relative to note length. This provides a general picture where males seasonally increase song effort and decrease trill rates and males with higher song effort and slower trills (i.e., longer notes) have larger clutches.

Regarding voiced modulation score, we found it declined as the fertile window approached while unvoiced modulation score did not change seasonally. Neither explained variation in reproductive success, unlike what would be expected for a signal of quality (Hau et al. 2017). Unvoiced modulation score was significantly higher during agonistic interactions, while otherwise both scores were unchanged across context. However, the increase in unvoiced modulation score during song duels was entirely explained by decreased gap length. In fact, frequency travelled and frequency jumps *per se* did not change with context and showed no biologically relevant variation associated with fertility or reproductive success. At first, these results appear in contrast with previous studies on different species that suggest that vocal performance is reflected in higher frequency modulation skill. However, close inspection reveals that our results do not contradict previous studies. Bosca et al. (2025) found that unvoiced modulation score increased during simulated intrusions, but, as in blue tits, frequency jumps were unchanged and only the gaps between notes were shortened. Most playback studies investigating vocal performance manipulated song by shortening gaps, or selecting notes of varying length (Ballentine et al. 2004; Drăgănoiu et al. 2002; Goodwin and Podos 2014; Illes et al. 2006; Phillips and Derryberry 2017a; Phillips and Derryberry 2017b). These manipulations would result in higher unvoiced modulation score (i.e., lower vocal deviation) but song discrimination can be explained entirely by differences in note length or in gap length. Furthermore, our results are consistent with a recent finding where the extent of frequency modulation and trill rates can be determined by biogeographic patterns and not by sexual selection (Bacquelé et al. 2026).

## Conclusion

Our results reveal a phenotypic boundary between song length and note length in blue tits, perhaps reflecting a respiratory constraint. Song effort score, which measures how close birds sing to this boundary, peaked during the fertile period and predicted clutch size in males, supporting it as a fitness indicator in birdsong. Song length alone, however, was a poor predictor of reproductive success, because its value as a signal only emerged relative to the note length that constrains it. Regarding behavioural context, singing duels were characterised by lower song effort scores whereas alarm songs had higher scores. Nevertheless, both contexts showed faster trills than solo songs, supporting trill rate as a motivation or arousal indicator. From acoustic to visual or vibrational modalities, trade-offs affecting motor performance are common in animal displays. Describing these performance challenges and how display components interact will improve the quantification of inter-individual variation on which selection acts. Revisiting previous studies in this light may resolve apparently conflicting results and help unify our understanding of the role of motor performance in animal communication across modalities and taxa.

