## Supplementary material for "A new fitness indicator in birdsong: a trade-off between song length and note length"

**Supplementary Methods**

*­­	Estimation of boundary trade-offs*

*Reciprocal quantile regressions.*

For each trade-off, both traits were z-standardized across both sexes (mean-centred and divided by their standard deviation), and two mixed-effects quantile regressions were fitted with the *lqmm* function, one with each trait as the response. Each model included the predictor, sex and their interaction, and individual identity as a random intercept to account for repeated measures. Because of this random intercept, the fitted quantiles describe the boundary of an average individual rather than the edge of the pooled distribution. For the song length–note length trade-off, an upper boundary, both reciprocal models were fitted at the 90% quantile. For the other three trade-offs, lower boundaries, the models with gap length ~ note length, note length ~ frequency travelled and gap length ~ frequency jump were fitted at the 10% quantile, and the reciprocal models at 90% quantile. Sex-specific lines were obtained from the coefficients of each model (Table S1; Figure S1).

*Boundary and performance scores.*

The boundary trade-off for each sex was the bisector of the two reciprocal lines, calculated in the standardized space, so that both traits weighed equally in the distance to the boundary whatever their original units. Boundaries were back-transformed to the original units only for plotting (Figures 2, S1–S2). For each observation, the performance score was the signed orthogonal distance to the bisector, in standardized units, with positive values beyond the boundary. Because the bisector halves the angle between the two reciprocal lines, this distance is proportional to the sum of the signed orthogonal distances to both lines, the consensus measure of the double quantile regression approach (Cardoso 2019). We confirmed this empirically: across all male observations, both measures were virtually identical (r ≥ 0.998; Figure S1).

*Confidence intervals and sex differences.*

Confidence intervals for the boundaries were obtained by a paired bootstrap (300 replicates): individuals were resampled, both reciprocal models were refitted on the same resampled individuals, and the bisector was recalculated for each sex. A trade-off was considered present when the 95% percentile interval of the boundary slope did not include zero, and a sex difference in the slope or intercept of the boundary was considered significant when the 95% percentile interval of the male − female difference did not include zero (Table S1). Sex differences in each reciprocal model were assessed from the sex and interaction terms of the quantile regressions. R code is provided in the data repository.

**Song and fitness**

All models described below were generalized Poisson GLMMs of clutch size including laying date and partner age as fixed effects and male identity as a random intercept, as in the main model, and were fitted on the same data set, so that their AICc values were directly comparable. All continuous predictors were scaled.

*Single-score models*

The averaged model (Table 2) estimates partial effects, that is, the effect of each score when the others are held constant. To obtain the effect of each score on its own, we fitted four models, each including a single score as the only song predictor. P values were corrected for multiple testing with the Holm method, and each model was compared by AICc with a model including only the covariates (Table S6).

Performance scores vs. raw traits. Each score is derived from two raw traits, so its association with clutch size could simply reflect one of them. We compared by AICc six models with the following song predictors: (i) none (covariates only); (ii) song effort score; (iii) song length; (iv) song length and note length, the two traits that define song effort score; (v) the four performance scores; and (vi) the five raw traits that constitute the four trade-offs (song length, note length, gap length, frequency travelled and frequency jump) (Table S7; Figure S4).

*Robustness to repertoire composition.*

Song types are largely defined by the song traits used here, so similar songs could be assigned to different song types if one male sings them with longer notes, shorter gaps or longer trills. Repertoire composition could therefore confound the association between performance and clutch size. To reduce potential bias in song-type categorization associated with performance level, we grouped the 14 song types in our population into five song-type clusters, and excluded clusters sung by fewer than three males. In the main text, we refer to these clusters as song types. To test whether song effort score reflects individual capacity regardless of song-type categorization, we split each male’s mean song effort score into two components (Van De Pol and Wright, 2009). For each song-type cluster, the population mean was calculated as the mean of the male means within the cluster, so that each male contributed equally regardless of the number of songs recorded. For each song, the between-cluster component was the population mean of its cluster and the within-cluster component was the deviation of the song from that mean; both were averaged per recording day and per breeding attempt in the same way as the scores, so that they add up to the male’s mean value. Both components were entered together, in place of the score, in a model with the covariates (Table S8A). A metric that reflects general individual capacity should be associated with clutch size within clusters: whatever the song type recorded, higher-quality males should sing it closer to the boundary. A between-cluster association would instead indicate that higher-quality males preferentially sing song types that lie closer to the boundary, so that repertoire composition itself carries information on quality. We applied the same decomposition to raw song length (Table S8B; Figure S6).

*Note–gap axes.*

Note length and gap length lengthen together, particularly across song types, which makes their separate effects, and the partial effect of recovery score hard to interpret. We therefore standardized both traits across breeding attempts (z-scores) and re-expressed them as two uncorrelated axes:

$$\text{Tempo}=\frac{z_{N}+z_{G}}{\sqrt{2}},\quad\quad\text{Contrast}=\frac{z_{N}-z_{G}}{\sqrt{2}}$$

where z_N_ and z_G_ are the standardized note length and gap length. Tempo describes how slow a male’s trills were (notes and gaps both long), and contrast how much of each note–gap cycle was filled with sound (notes long relative to their gaps, analogous to recovery score). These are the principal components of the two standardized traits and are uncorrelated by construction. Tempo and contrast were entered together, in place of note length and gap length, in a model that also included song length and the covariates (Table S8C).

*Model diagnostics.*

Residuals of the main model were checked with simulated quantile residuals (‘DHARMa’), including tests for uniformity (Kolmogorov–Smirnov), dispersion and outliers (Figure S3). All variance inflation factors were ≤ 3. Models were considered converged when the Hessian matrix was positive definite.

**Supplementary figures**

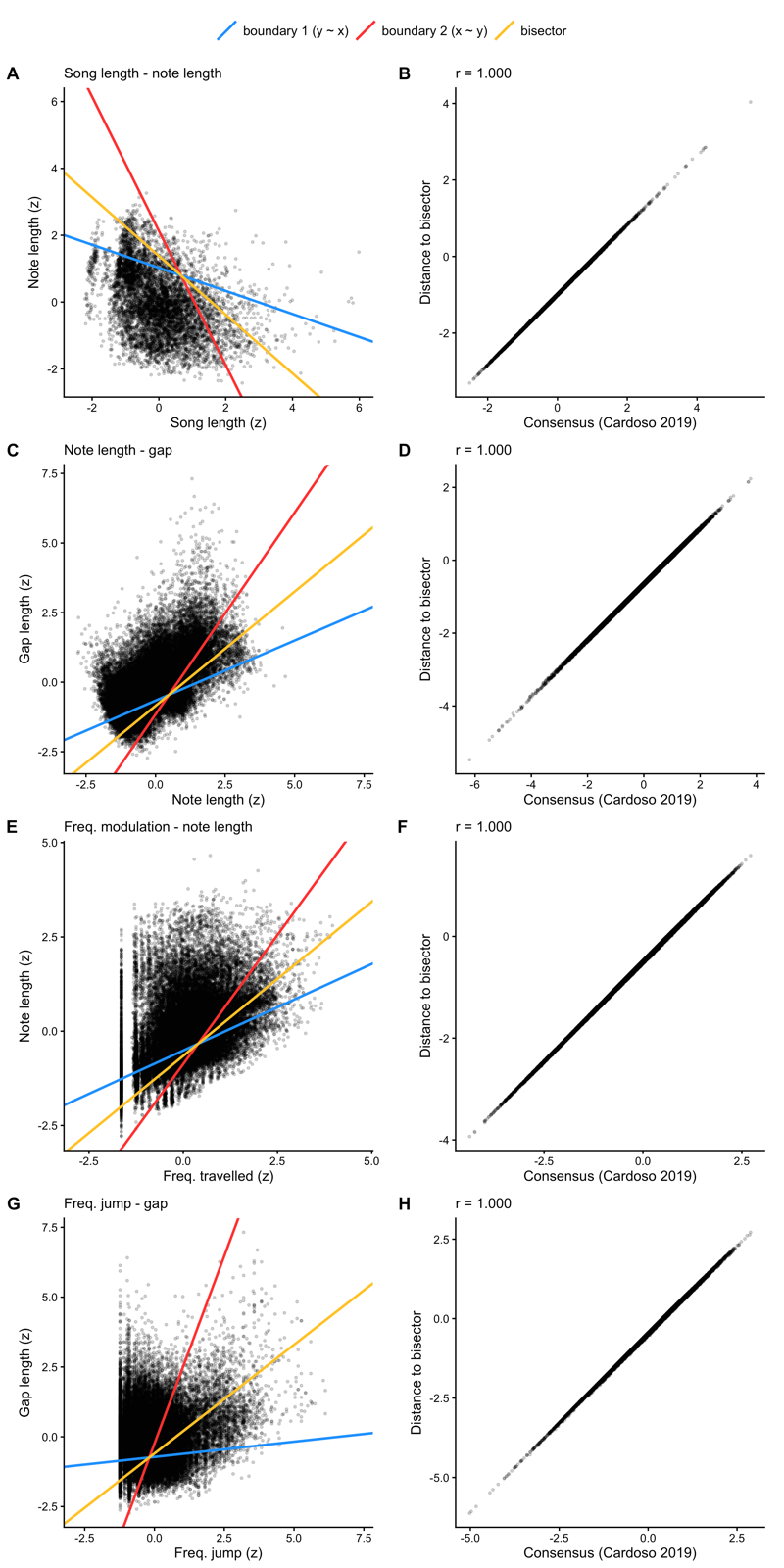

Figure S1 – Reciprocal quantile regressions and comparison of the distance to the bisector with the consensus metric of Cardoso (2019), in males. Left: phenotypic space (z-standardized traits) described by (A) note length and song length, (B) gap length and note length, (C) note length and frequency travelled and (D) gap length and frequency jump. Blue and red lines are the two reciprocal mixed-effects quantile regressions (boundary 1: y ~ x; boundary 2: x ~ y), and the yellow line is their bisector, which defines the boundary shown in Figure 2 (Table S1). Right: distance of each observation to the bisector against the consensus metric (mean of the standardized residuals of both quantile regressions); r, Pearson correlation.

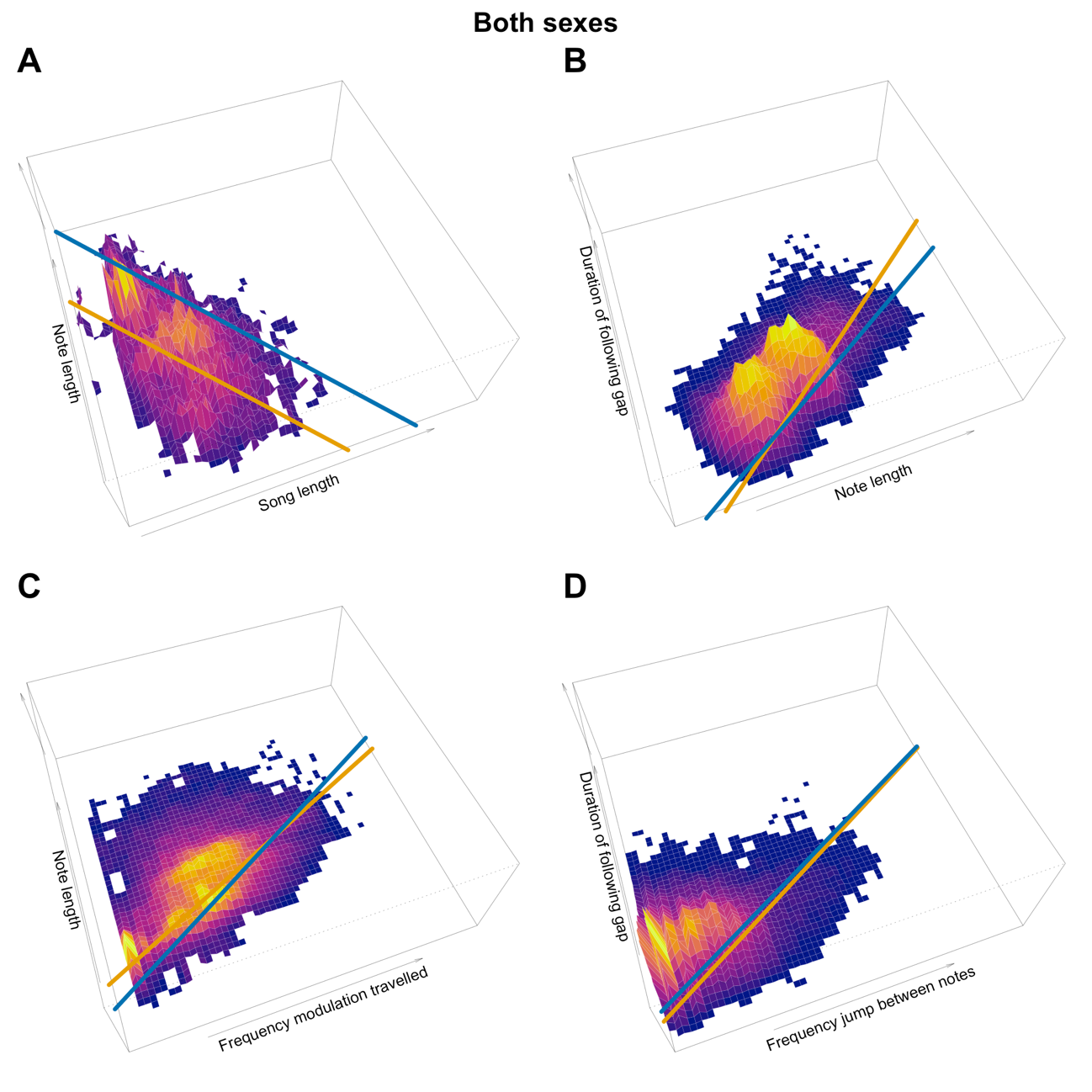

Figure S2 – Density of observations in the phenotypic space. Height and colour of each bin show the relative number of observations. Boundaries are represented for each sex, with the consensus boundary estimated for males (blue) and females (yellow) from a single model including sex and its interaction with the predictor (Table S1).

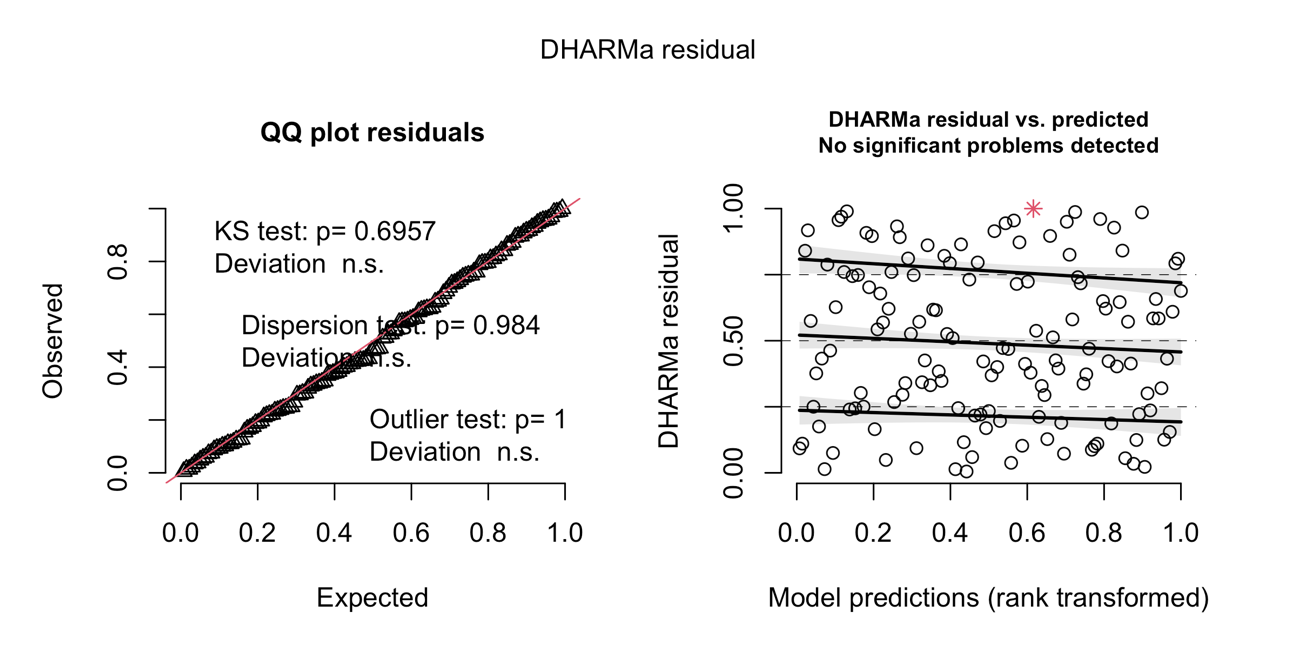

Figure S3 – Residual diagnostics of the main clutch size model. Simulated quantile residuals (DHARMa) of the full generalized Poisson GLMM. Left: QQ plot against the uniform distribution, with Kolmogorov–Smirnov, dispersion and outlier tests. Right: residuals against model predictions, with quantile regressions at 0.25, 0.5 and 0.75.

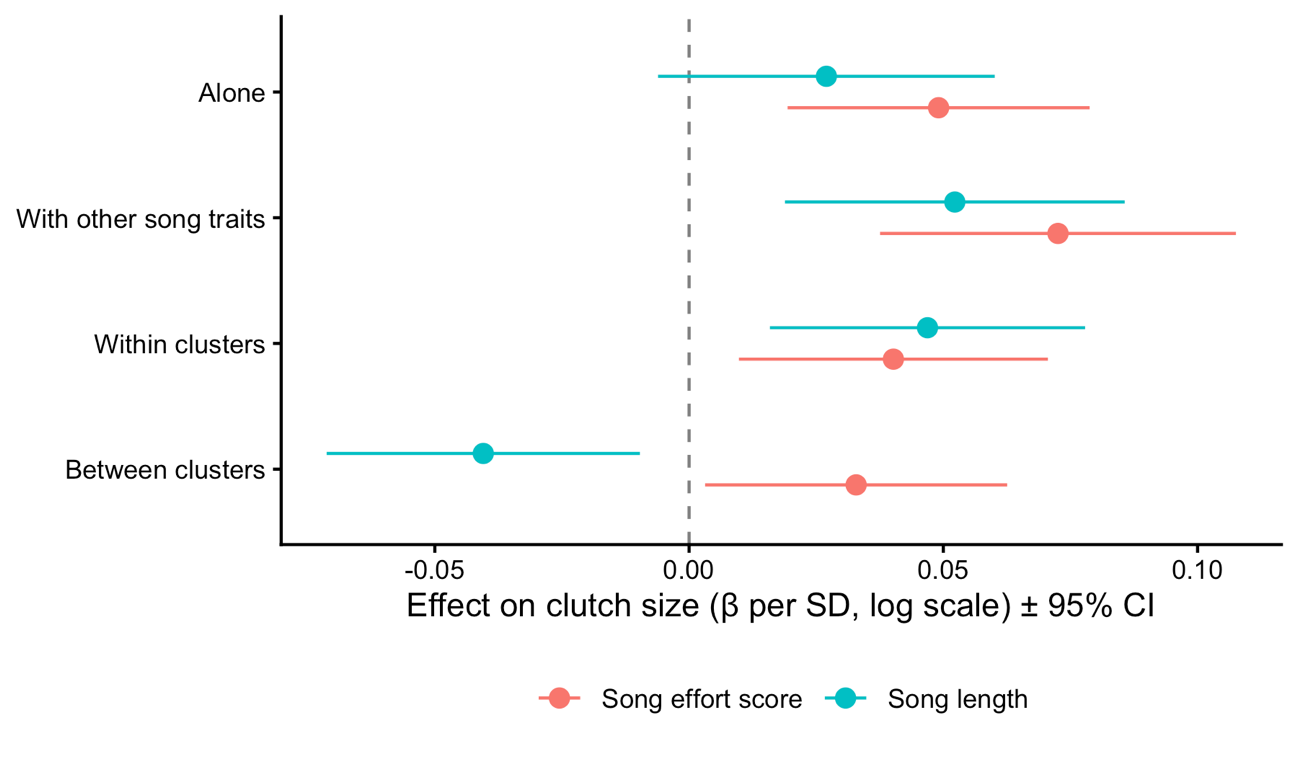

Figure S4 – Song effort score vs. song length as predictors of clutch size. Effect on clutch size (β per SD, log scale, ± 95% CI) of song effort score and raw song length when fitted alone with the covariates (Tables S6-S7), together with the other song traits (the four scores or the five raw traits; Table S7), and split into within- and between-cluster components (Table S8A–B).

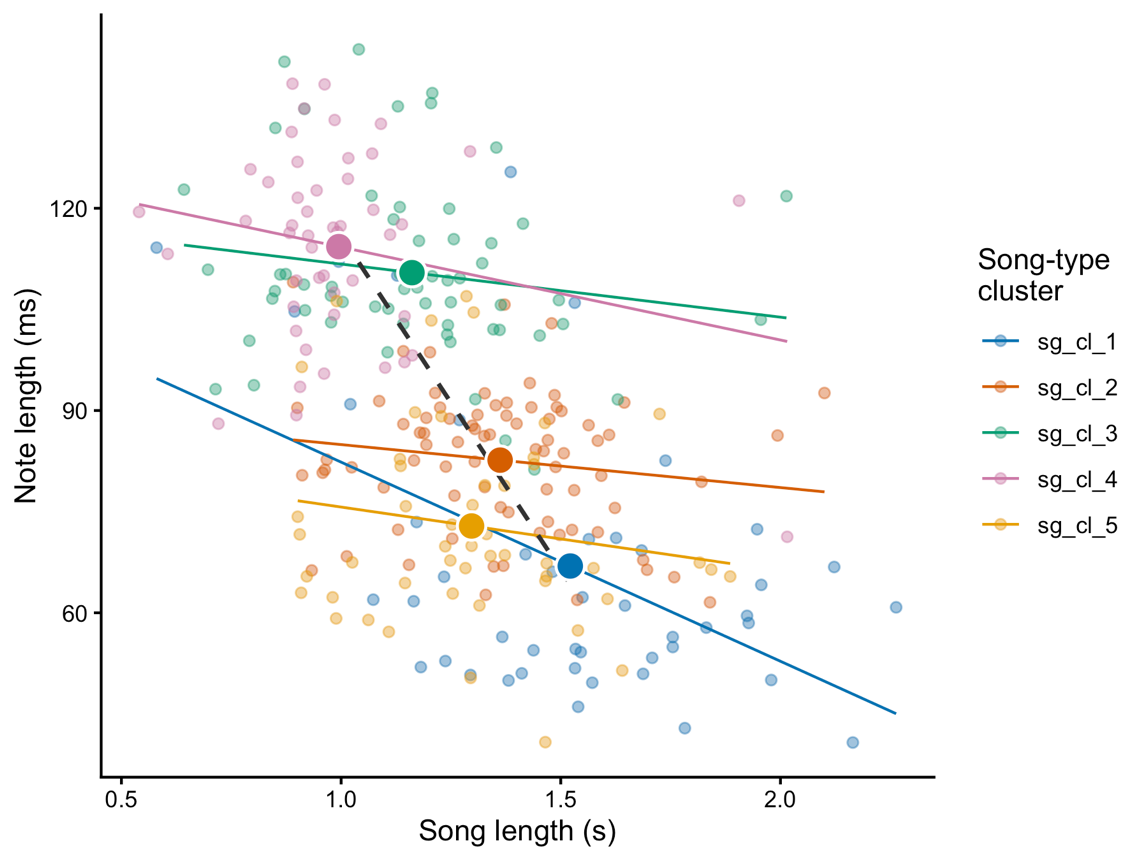

Figure S5 – Song length and note length across song-type clusters. Small points are the mean of one male within one song-type cluster; large points are cluster means (mean of male means). Thin lines show the relationship within each cluster and the dashed line the relationship between cluster means. Only males included in the fitness analysis and clusters sung by at least three males are shown.

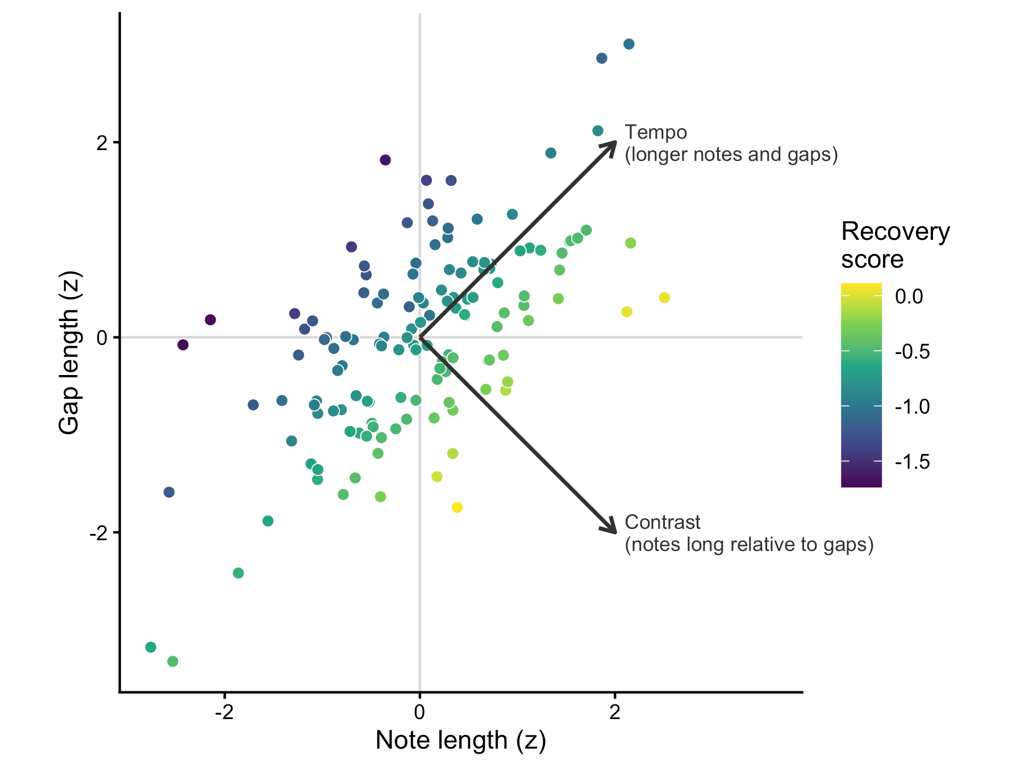

Figure S6 – Note length and gap length as two uncorrelated axes. Mean note length and gap length per breeding attempt (standardized), coloured by recovery score. Arrows show the tempo axis (longer notes and longer gaps) and the contrast axis (notes long relative to their gaps) used in Table S8C.

| **Table S1 – Boundary trade-offs in blue tit song by sex** | | | | | | | | |
| --- | --- | --- | --- | --- | --- | --- | --- | --- |
|  |  |  | **Consensus** |  | **y ~ x** |  | **x ~ y** |  |
| **Trait (y)** | **Trait (x)** | **Sex** | **Intercept [95%CI]** | **Slope [95%CI]** | **Intercept [95%CI]** | **Slope [95%CI]** | **Intercept [95%CI]** | **Slope [95%CI]** |
| Note length | Song length | Female | 0.146 [0.043,0.174] | -0.051 [-0.065,0.048] | 0.093 [0.074,0.108] | -0.004 [-0.016,0.005] | 1.48 [1.204,1.911] | -4.086 [-8.211,0.475] |
| N = 7864 songs |  | Male | 0.187 [0.166,0.194] | -0.055 [-0.06,-0.049] | 0.138 [0.12,0.148] | -0.021 [-0.029,-0.015] | 2.337 [2.075,2.449] | -8.006 [-9.478,-6.415] |
| Inter-note gap length | Note length | Female | -0.019 [-0.025,0.099] | 0.711 [-0.754,0.804] | 0.03 [0.021,0.05] | 0.131 [-0.189,0.215] | 0.075 [0.044,0.089] | 0.21 [-0.082,0.589] |
| N = 37298 notes |  | Male | -0.009 [-0.013,-0.002] | 0.555 [0.511,0.601] | 0.015 [0.008,0.02] | 0.292 [0.231,0.322] | 0.047 [0.037,0.055] | 1.02 [0.875,1.1] |
| Note length | Freq. modulation | Female | 0.047 [0.029,0.056] | 0.318 [0.278,0.534] | 0.062 [0.035,0.074] | 0.093 [0.05,0.355] | -0.031 [-0.071,0.026] | 1.404 [0.753,1.925] |
| N = 35016 notes |  | Male | 0.036 [0.029,0.047] | 0.395 [0.345,0.44] | 0.052 [0.048,0.066] | 0.222 [0.171,0.244] | -0.018 [-0.059,0.017] | 1.511 [1.06,1.863] |
| Inter-note gap length | Freq. jump | Female | 0.021 [0.013,0.033] | 0.437 [0.351,0.602] | 0.031 [0.026,0.044] | 0.171 [0.031,0.249] | -0.002 [-0.029,0.033] | 1.052 [0.319,1.338] |
| N = 34588 notes |  | Male | 0.025 [0.015,0.03] | 0.421 [0.393,0.47] | 0.035 [0.03,0.043] | 0.059 [0.005,0.106] | 0.003 [-0.006,0.028] | 0.689 [0.362,0.891] |
| Consensus boundary (bisector of the two reciprocal quantile regression lines) and the two reciprocal mixed-effects quantile regressions (y ~ x and x ~ y) for each trade-off, with sex and its interaction with the predictor in each model. Models were fitted on z-standardized traits and lines were back-transformed to the original units; estimates of the x ~ y models are expressed in y ~ x units. Values are estimates with bootstrap 95% CIs. N = number of songs (song length–note length) or notes (other trade-offs). | | | | | | | | |

**Supplementary table**

| **Table S2 – Seasonal variation of male song: smooth terms of the GAMMs.** | | | | | |
| --- | --- | --- | --- | --- | --- |
| **NON-PARAMETRIC** |  |  |  |  |  |
| **Response variable** | **Predictors** | **EDF** | **Ref. df** | **F** | **P** |
| Song effort score | Weeks to first egg | 3.695 | 3.942 | 20.427 | < 0.001 |
|  | ID | 0.001 | 67 | 0 | 1 |
| Recovery score | Weeks to first egg | 3.795 | 3.973 | 12.577 | < 0.001 |
|  | ID | 0.001 | 67 | 0 | 1 |
| Voiced mod. score | Weeks to first egg | 2.237 | 2.731 | 18.78 | < 0.001 |
|  | ID | 0 | 67 | 0 | 1 |
| Unvoiced mod. score | Weeks to first egg | 3.443 | 3.82 | 1.482 | 0.128 |
|  | ID | 0.001 | 67 | 0 | 1 |
| Song length | Weeks to first egg | 3.633 | 3.918 | 11.686 | < 0.001 |
|  | ID | 0.001 | 67 | 0 | 1 |
| Note length | Weeks to first egg | 3.696 | 3.943 | 25.208 | < 0.001 |
|  | ID | 0.001 | 67 | 0 | 1 |
| Gap length | Weeks to first egg | 3.672 | 3.934 | 5.038 | < 0.001 |
|  | ID | 0.001 | 67 | 0 | 1 |
| Freq. modulation travelled | Weeks to first egg | 1 | 1 | 3.599 | 0.058 |
|  | ID | 0 | 67 | 0 | 1 |
| Frequency jump | Weeks to first egg | 1 | 1 | 17.166 | < 0.001 |
|  | ID | 0.001 | 67 | 0 | 1 |
| Trill rate (notes/s) | Weeks to first egg | 2.071 | 2.546 | 15.374 | < 0.001 |
|  | ID | 0.001 | 67 | 0 | 1 |
| Effective degrees of freedom (EDF), reference degrees of freedom (Ref. df), F and P for the smooth of weeks to first egg and the random effect of individual, for the four performance scores, the raw traits that define them and trill rate. EDF > 1 indicates a non-linear seasonal change. EDF of individual is close to zero because all variables were standardized within individual and song type. Parametric terms are shown in Table 1 and Table S4 (raw traits). | | | | | |

| **Table S3 – Timing of the seasonal maximum and minimum of each song trait.** | | | | | |
| --- | --- | --- | --- | --- | --- |
| **Variable** | |  | Week | 2.5% CI | 97.5% CI |
| Song effort score | | Maximum | -0.402 | -1.107 | 2 |
|  | | Minimum | -14 | -14 | -6.569 |
| Recovery score | | Maximum | -0.691 | -1.363 | 2 |
|  | | Minimum | -14 | -14 | -14 |
| Voiced modulation score | | Maximum | -14 | -14 | -7.321 |
|  | | Minimum | 2 | 2 | 2 |
| Unvoiced modulation score | | Maximum | -10.797 | -14 | -1.684 |
|  | | Minimum | 2 | -14 | 2 |
| Song length | | Maximum | -1.219 | -1.684 | -0.113 |
|  | | Minimum | -14 | -14 | -6.697 |
| Note length | | Maximum | 2 | -0.115 | 2 |
|  | | Minimum | -14 | -14 | -6.248 |
| Inter-note gap length | | Maximum | -14 | -14 | -6.056 |
|  | | Minimum | -1.091 | -10.173 | 2 |
| Frequency travelled | | Maximum | 2 | -14 | 2 |
|  | | Minimum | -14 | -14 | 2 |
| Frequency jump | | Maximum | -14 | -14 | -14 |
|  | | Minimum | 2 | 2 | 2 |
| Trill rate | | Maximum | -14 | -14 | -7.337 |
|  | | Minimum | 2 | 2 | 2 |
| Week relative to first egg (0 = laying week) at which the GAMM prediction for solo song reached its maximum and minimum, with 95% CIs from 1,000 simulations from the multivariate normal distribution of the model coefficients. The sampled period spanned weeks −14 to +2. | | | | | |

| **Table S4 – Effect of behavioural context on the raw song traits and trill rate.** | | | | | | | |
| --- | --- | --- | --- | --- | --- | --- | --- |
| **PARAMETRIC** |  |  |  |  |  |  |  |
| **Response variable** | **Predictors** | **Estimate** | **SE** | **2.5% CI** | **97.5% CI** | ***t*** | **P** |
| Song length | Intercept | -0.024 | 0.034 | -0.091 | 0.042 | -0.719 | 0.473 |
|  | Agonistic vs. solo | -0.451 | 0.181 | -0.806 | -0.096 | -2.491 | 0.013 |
|  | Alarm vs. solo | 0.285 | 0.142 | 0.006 | 0.564 | 2.001 | 0.046 |
| Note length | Intercept | -0.017 | 0.033 | -0.081 | 0.046 | -0.532 | 0.595 |
|  | Agonistic vs. solo | -0.026 | 0.174 | -0.367 | 0.316 | -0.148 | 0.883 |
|  | Alarm vs. solo | -0.182 | 0.137 | -0.45 | 0.087 | -1.328 | 0.185 |
| Inter-note gap length | Intercept | 0.039 | 0.035 | -0.029 | 0.108 | 1.123 | 0.262 |
|  | Agonistic vs. solo | -0.386 | 0.187 | -0.753 | -0.019 | -2.059 | 0.04 |
|  | Alarm vs. solo | -0.304 | 0.147 | -0.592 | -0.015 | -2.063 | 0.04 |
| Freq. modulation | Intercept | 0.003 | 0.035 | -0.066 | 0.073 | 0.093 | 0.926 |
|  | Agonistic vs. solo | 0.075 | 0.19 | -0.296 | 0.447 | 0.397 | 0.691 |
|  | Alarm vs. solo | 0.058 | 0.147 | -0.231 | 0.347 | 0.394 | 0.694 |
| Freq. jump | Intercept | 0.012 | 0.035 | -0.058 | 0.081 | 0.331 | 0.741 |
|  | Agonistic vs. solo | -0.168 | 0.189 | -0.539 | 0.204 | -0.885 | 0.376 |
|  | Alarm vs. solo | 0.061 | 0.147 | -0.227 | 0.35 | 0.415 | 0.678 |
| Trill rate (notes/s) | Intercept | -0.012 | 0.034 | -0.078 | 0.055 | -0.342 | 0.733 |
|  | Agonistic vs. solo | 0.742 | 0.181 | 0.387 | 1.096 | 4.1 | < 0.001 |
|  | Alarm vs. solo | 0.353 | 0.142 | 0.075 | 0.631 | 2.487 | 0.013 |

Parametric terms of the GAMMs described in Table 1, fitted to the traits that define the four trade-offs and to trill rate. Variables were standardized within individual and song type; the intercept corresponds to solo song.

| **Table S5 – Model selection for clutch size.** | | | | | | | | | | | |
| --- | --- | --- | --- | --- | --- | --- | --- | --- | --- | --- | --- |
| Intercept | Partner age (older) | Laying date | Unvoiced modulation score | Voiced modulation score | Recovery score | Song effort score | df | logLik | AICc | ΔAICc | Weight |
| 2.187 | – | -0.082 | – | – | -0.052 | 0.070 | 6 | -252.64 | 517.92 | 0.00 | 0.230 |
| 2.164 | + | -0.084 | – | – | -0.046 | 0.067 | 7 | -251.65 | 518.17 | 0.25 | 0.203 |
| 2.186 | – | -0.081 | 0.018 | – | -0.061 | 0.076 | 7 | -252.07 | 519.00 | 1.08 | 0.134 |
| 2.162 | + | -0.083 | 0.019 | – | -0.055 | 0.074 | 8 | -251.04 | 519.21 | 1.29 | 0.121 |
| 2.187 | – | -0.082 | – | 0.010 | -0.052 | 0.074 | 7 | -252.46 | 519.78 | 1.86 | 0.091 |
| 2.163 | + | -0.084 | – | 0.012 | -0.045 | 0.072 | 8 | -251.41 | 519.94 | 2.02 | 0.084 |
| Generalized Poisson GLMMs including all combinations of the four song performance scores, laying date and partner age (male identity as random intercept), ranked by AICc. Only models within ΔAICc < 4 of the best model are shown; models within ΔAICc < 2 were averaged (Table 2). Values are coefficients on the log scale (continuous predictors scaled); "+" indicates that partner age was included and "–" that a term was absent. Akaike weights are relative to all models. | | | | | | | | | | | |

| **Table S6 – Models of clutch size variation fitted using a single performance score as predictor** | | | | | | |
| --- | --- | --- | --- | --- | --- | --- |
| **Song performance score** | **β [95% CI]** | **Change in clutch size per SD (%)** | **z** | **P** | **P (Holm)** | **ΔAICc vs. covariates only** |
| Song effort score | 0.049 [0.019, 0.079] | 5.0 | 3.24 | 0.001 | 0.005 | -7.85 |
| Recovery score | -0.017 [-0.049, 0.015] | -1.7 | -1.02 | 0.307 | 0.922 | 1.15 |
| Voiced modulation score | -0.012 [-0.044, 0.019] | -1.2 | -0.77 | 0.442 | 0.922 | 1.60 |
| Unvoiced modulation score | -0.010 [-0.042, 0.022] | -1.0 | -0.62 | 0.536 | 0.922 | 1.81 |
| Association between each song performance score and clutch size when fitted as the only song predictor, in generalized Poisson GLMMs with laying date, partner age and male identity as random intercept. Estimates are on the log scale per SD; the change in clutch size per SD is exp(β) − 1. P values are corrected for the four tests with the Holm method. ΔAICc is relative to the covariates-only model; negative values indicate a better fit. N = 138 breeding attempts from 96 males. | | | | | | |

| **Table S7 – Song performance scores vs. raw traits.** | | | | | | | |
| --- | --- | --- | --- | --- | --- | --- | --- |
| **Model** | **Song-length term** | **β [95% CI]** | **P** | **K** | **AICc** | **ΔAICc vs. covariates only** | **ΔAICc vs. best** |
| Covariates only | - | - | - | 5 | 530.88 | 0.00 | 9.42 |
| Song effort score | Song effort score | 0.049 [0.019, 0.079] | 0.001 | 6 | 523.03 | TRUE | -7.85 |
| Song length | Song length | 0.027 [-0.006, 0.060] | 0.110 | 6 | 530.51 | TRUE | -0.37 |
| Song length + note length | Song length | 0.045 [0.010, 0.079] | 0.011 | 7 | 525.31 | TRUE | -5.57 |
| Four scores (main model) | Song effort score | 0.073 [0.038, 0.108] | < 0.001 | 9 | 521.46 | TRUE | -9.42 |
| Five raw traits | Song length | 0.052 [0.019, 0.086] | 0.002 | 10 | 521.71 | TRUE | -9.17 |
| Six generalized Poisson GLMMs of clutch size fitted on the same breeding attempts, all including laying date, partner age and male identity as random intercept. The song length term of each model (song effort score or song length) is shown with its 95% CI. K, number of estimated parameters. ΔAICc is given relative to the covariates-only model and to the best model. “Five raw traits” = song length, note length, gap length, frequency travelled and frequency jump, the traits that define the four trade-offs. | | | | | | | |

| **Table S8 – Robustness of the association with clutch size to repertoire composition and to the covariation of note and gap length.** | | | | | |
| --- | --- | --- | --- | --- | --- |
| **Model** | **Predictor** | **β [95% CI]** | **SE** | **z** | **P** |
| A. Song effort score within vs. | Intercept | 2.152 [2.107, 2.198] | 0.023 | 92.55 | < 0.001 |
| between song-type clusters | Song effort score, within song-type cluster | 0.040 [0.010, 0.071] | 0.016 | 2.59 | 0.010 |
|  | Song effort score, between song-type clusters | 0.033 [0.003, 0.063] | 0.015 | 2.17 | 0.030 |
|  | Laying date | -0.085 [-0.110, -0.059] | 0.013 | -6.39 | < 0.001 |
|  | Partner age (older vs. first year) | 0.062 [0.005, 0.119] | 0.029 | 2.14 | 0.032 |
| B. Song length within vs. | Intercept | 2.158 [2.112, 2.203] | 0.023 | 93.11 | < 0.001 |
| between song-type clusters | Song length, within song-type cluster | 0.047 [0.016, 0.078] | 0.016 | 2.96 | 0.003 |
|  | Song length, between song-type clusters | -0.040 [-0.071, -0.010] | 0.016 | -2.57 | 0.010 |
|  | Laying date | -0.086 [-0.112, -0.060] | 0.013 | -6.50 | < 0.001 |
|  | Partner age (older vs. first year) | 0.053 [-0.004, 0.110] | 0.029 | 1.81 | 0.070 |
| C. Song length + note–gap axes | Intercept | 2.166 [2.120, 2.212] | 0.023 | 92.36 | < 0.001 |
|  | Song length | 0.053 [0.019, 0.086] | 0.017 | 3.06 | 0.002 |
|  | Tempo (longer notes and gaps) | 0.062 [0.031, 0.094] | 0.016 | 3.85 | < 0.001 |
|  | Contrast (notes long relative to gaps) | -0.020 [-0.050, 0.010] | 0.015 | -1.30 | 0.194 |
|  | Laying date | -0.083 [-0.108, -0.057] | 0.013 | -6.32 | < 0.001 |
|  | Partner age (older vs. first year) | 0.038 [-0.021, 0.096] | 0.030 | 1.27 | 0.205 |
| Generalized Poisson GLMMs of clutch size, all including laying date, partner age and male identity as random intercept. (A) Song effort score split into a within-cluster component (the male’s mean deviation from the population mean of his song-type clusters) and a between-cluster component (the mean population score of the clusters recorded for the male). (B) The same decomposition for raw song length. (C) Song length together with tempo (longer notes and longer gaps) and contrast (notes long relative to their gaps), two uncorrelated axes derived from note length and gap length. See Supplementary Methods. | | | | | |
